# EditR: A novel base editing quantification software using Sanger sequencing

**DOI:** 10.1101/213496

**Authors:** Mitchell G. Kluesner, Derek A. Nedveck, Walker S. Lahr, John R. Garbe, Juan E. Abrahante, Beau R. Webber, Branden S. Moriarity

## Abstract

CRISPR/Cas9-Cytidine deaminase fusion enzymes - termed Base Editors – allow targeted editing of genomic deoxcytidine to deoxthymidine (C→T) without the need for double stranded break induction. Base editors represent a paradigm-shift in gene editing technology, due to their unprecedented efficiency to mediate targeted, single-base conversion; however, current analysis of base editing outcomes rely on methods that are either imprecise or expensive and time consuming. To overcome these limitations, we developed a simple, cost effective, and accurate program to measure base editing efficiency from fluorescence-based Sanger sequencing, termed EditR. We provide EditR as a free online tool or downloadable desktop application requiring a single Sanger sequencing file and guide RNA sequence (baseeditr.com). EditR is more accurate than enzymatic assays, and provides added insight to the position, type and efficiency of base editing. Collectively, we demonstrate that EditR is a robust, inexpensive tool that will facilitate the broad application of base editing technology, thereby fostering further innovation in this burgeoning field.

## INTRODUCTION

Recently, several research groups have developed Cas9-Cytidine deaminase fusion enzymes for the purpose of gene editing with single base resolution (1–4). These base editors rely on the programmable specificity of the Cas9-guide RNA (gRNA) complex to localize a mutagenic cytidine deaminase enzyme to produce targeted deoxycytidine to deoxyuridine (C → U) mutations. Through DNA replication, deoxyuridine behaves like deoxythymidine resulting in C → T mutations (antisense G → A). By leveraging disparate outcomes in DNA repair, some base editors preferentially induce C → T mutations (canonical mutations) (1, 2), while others were developed for random mutagenesis (non-canonical) of C → T, G or A (antisense G → A, C or T) (3, 4). The single-nucleotide level resolution of base editing shows promise in gene therapy (5), agricultural engineering (6, 7) and basic scientific research (8, 9). Employing base editing in any laboratory setting requires the ability to quantify the efficiency, precision, accuracy and reproducibility of base editing. Demonstrably, all work published on base editing to date includes a quantitative assessment of base editing efficiency (1–13).

Of the several documented approaches to measuring base editing efficiency, all are limited in their precision, cost or delay in receiving results. Rapid and cost-effective approaches to measuring base editing are enzymatic cleavage assays such as the Cel I, T7E, Surveyor, or Guide-it Resolvase assays (14, 15). However, these assays are unable to discern the exact position and type of mutation because they only rely on the presence of a mismatch bubble formed in heteroduplexes of stochastically annealed DNA (16). This approach is suboptimal for base editing where adjacent Cs may be edited or non-canonical C → T, G or A mutations may occur, neither of which can be distinguished by enzymatic cleavage assay. As an alternative, bacterial colony sequencing of subcloned PCR amplicons can elucidate the specific outcomes of base editing (2, 7), but it is a time-consuming, laborious, and costly approach, making it impractical for mid- to high-throughput research. In comparison, the most informative method to measure base editing is next-generation deep sequencing (NGS) of the edited site (3, 4, 10, 11), but, this is the most expensive and time-consuming method and requires bioinformatics expertise.

In the analysis of insertion-deletion mutations (indels) from CRISPR/Cas9 editing, bioinformatic approaches using fluorescent capillary Sanger sequencing provide rapid and affordable methods to measure and characterize editing efficiency, most notably with the free web tools, Tracking of Indels by DEcomposition (TIDE) (https://tide.nki.nl/) and Poly Peak Parser (http://yosttools.genetics.utah.edu/PolyPeakParser/) (16, 17). These programs analyze secondary Sanger sequencing traces to delineate the composition and frequency of indel mutations. Inspired by these programs, we developed an accurate, fast, and low-cost method for the identification and quantification of base editing from fluorescent Sanger sequencing data. We provide this program, **EditR** (Edit deconvolution by inference of traces in R) as a free web tool (baseEditR.com) or an open source R Shiny application that can run on a local desktop. EditR requires only a single Sanger sequencing file of a base edited sample and the sequence of the gRNA protospacer to disentangle the outcomes of base editing.

## MATERIALS AND METHODS

### Plasmids and gRNA design

The identity of all plasmids in this study were confirmed by Sanger sequencing and restriction digests followed by gel electrophoresis. All base editing was carried out using pCMV-BE3 developed by Dr. David Liu’s Lab (Addgene # 73021, 1). BE3 was the first published base editor and has arguably the most comprehensive examination of activity *in vitro* and in cell culture. Guide RNAs (gRNAs) for use with BE3 were designed to the loci of interest using parameters outlined in previous publications including size of the editing window, identity of preceding base, distance from the protosopacer adjacent motif (PAM), and PAM specificity (1) (Supplementary Table S1). gRNAs were ordered as complementary oligonucleotides: 5’-CACC-G-protospacer-3’ and 5’-AAAC-reverse complement protospacer-C-3’ (Integrated DNA Technologies, IDT). Complementary oligonucleotides were annealed and phosphorylated with T4 PNK (NEB) and 10x T4 ligation buffer (NEB) in a thermocycler using the protocol: 37°C – 30 minutes, 95°C – 5 minutes, step-down to 25°C at 5°C/min. pENTR221-U6 stuffer vector was digested with BsmBI restriction enzyme, FastAP alkaline phosphatase (Fermentas), and 10x Tango Buffer overnight at 37°C. Linearized pENTR221-U6 and 1:200 diluted annealed and phosphorylated oligonucleotides were ligated together with T4 DNA ligase and buffer (NEB) at room temperature for ≥1 hour. Ligation reactions were transformed into DH10β *E. coli* (ThermoFisher Scientific), plated for single colonies, picked, cultured overnight, and plasmid DNA was extracted with a proprietary kit (GeneJET Plasmid Miniprep Kit, ThermoFisher Scientific). Plasmid identity was dually confirmed with HindIII-Hifi and PvuII-Hifi (NEB) restriction digest-gel electrophoresis and Sanger sequencing of gRNA region (ACGT, Inc). Confirmed plasmids were re-transformed, and plasmid DNA was extracted with a proprietary kit (HiSpeed Plasmid Maxi Kit, Qiagen).

### Cell line culturing and transfection

Cell lines were maintained at 37°C, 5% CO_2_ under 80% confluency and passaged 1:10 three times per week. HCT116 cells were maintained in DMEM (ThermoFisher Scientific) and HOS cells were maintained in EMEM (ATCC). All cell culturing media was supplemented with 10% FBS and 1x Penicillin-streptomycin. Puromycin selection was performed using media containing 1 μg/mL puromycin. HCT116 and HOS cells ≤ 80% confluent were electroporated using 1 μg pENTR221-gRNA, 1 μg pCMV-BE3, and 500 ng pmaxGFP (Lonza) according to the manufacturer’s protocol (Neon Transfection System, Life Technologies), and plated into a polylysine-coated 6-well plate. Twenty-four hours post electroporation percent GFP^+^ cells were observed to qualitatively assess transfection efficiency and genomic DNA was harvested 72 hours post electroporation.

### Co-Transposition and single colony isolation

Co-Transposition was performed via electroporation of an additional 500ng of PB-CG-Luciferase-EGFP (Puro) *PiggyBac* transposon and 500ng of hyperactive *PiggyBac* transposase as previously described (15) alongside the aforementioned pCMV-BE3 and pENTR221-gRNA plasmids. In principle, cells that obtain a transposition event integrating the puromycin resistance gene are also more likely to have taken up Cas9/BE3 and gRNA expression plasmids, thus more likely to be edited. Twenty-four hours post electroporation percent GFP^+^ cells were observed to qualitatively assess transfection and genomic DNA was harvested from half of the cells 72 hours post electroporation. The remaining cells were plated with puromycin supplemented media for single colony isolation on a 15-cm polylysine coated dish or serially diluted in 96-well-plate. Single colonies on 15-cm plates were allowed to grow for 14 days, or until visible to the naked eye, and isolated with colony isolators and Trypsin-EDTA (Fisher), or picked with a 10 μL pipette tip of Trypsin-EDTA and transferred to a 24-well dish. Once clones reached >90% confluency genomic DNA was harvested to assess editing.

### Cel I surveyor nuclease assay

Primers were designed to produce amplicons approximately 300-400 bp in length with the target site off-centered in the amplicon. Genomic DNA was PCR amplified with AccuPrime Taq DNA Polymerase, high fidelity (Invitrogen), 10X Accuprime buffer, and 5% DMSO, and ran on 1% agarose gel-electrophoresis and gel extracted (QiaQuick Gel Extraction Kit, Qiagen) or PCR purified (PCR Purification Kit, Qiagen). PCR products were denatured and annealed in a thermocycler using the <u>manufacturer’s protocol (IDT)</u>. Three microliters of denatured PCR products were combined with 1 μL 1x AccuPrime buffer II (Thermofisher), 0.7 μL of surveyor nuclease, and 0.7 μL of surveyor enhancer (IDT) then incubated at 42°C for 20 minutes. Reactions were terminated with Ficoll loading dye and ran on an agarose gel (2% m/v, 0.06 μL/mL ethidium bromide) in TAE buffer or a polyacrylamide gel in TBE buffer. Gel was imaged and the fraction of amplicons edited was quantified in ImageJ with the formula *F*_*Edited*_ = (*b + c*) / (*a + b + c*), where *a* is the integrated intensity of the undigested PCR band and *b* and *c* are the integrated intensities of each band as previously described (14).

### Sanger sequencing

Purified PCR product (1 ng/μL), primer (20 pmol/μL) and Big Dye Terminator v3.1 (4 μL) were brought to 12μL in molecular H_2_O and sequenced using the protocol: 95°C – 1 min [95°C – 30 sec, 56°C – 30 sec, 60°C – 1 min] x 24, 16°C – hold. Sequencing reactions were analyzed on an Applied Biosystems 3730 DNA Analyzer.

### Illumina Next-Generation Deep Sequencing

Primers were designed using Primer3 and Primer-BLAST to 300-500 bp regions of interest, with Nextera universal adaptors flanking the site-specific primer (Supplementary Table S1). Genomic DNA was PCR amplified in one step using AccuPrime Taq DNA Polymerase, high fidelity according to the manufacturer’s protocol (Invitrogen). Samples were submitted to the University of Minnesota Genomics Center for subsequent amplification with indexed primers and sequencing on a MiSeq 2×300 bp run (Illumina). A minimum of 1000 read-pairs were generated per sample.

Sequencing reads were demultiplexed using bcl2fastq2 (Illumina). FastQC (version 0.11.5) (18) was used to assess the quality of the data. Overlapping read-pairs were assembled with Pear (version 0.9.10) (19). Non-overlapping read-pairs and read-pairs with an assembled length 5 bp longer or shorter than the length of the amplicon reference sequence were discarded. Needle (EMBOSS version 6.5.7) (20) was used to generate optimal global sequence alignments between each assembled read and the amplicon reference sequence. The numbers of insertions, deletions, and substitutions at each base of the reference amplicon sequence were counted. Alignments of the 10 most common amplicon reads were visualized using MView (version 1.52) (21).

### EditR Software Development

To determine if the measured percent editing was significant, we implemented a null-hypothesis significance testing approach using a null-distribution modeled from the background noise. The null distribution is generated by trimming the first 20 bases of the sequence and removing the 20 bases of the protospacer. Additionally, bases that fall within the 10th percentile of total area are removed as small peaks are associated with poor initial primer binding and poor end extension (22). To account for the variability in sequencing, the user can manually select the region to model the null distribution in case the default trimming doesn’t effectively remove low quality sequencing. Next, the value of every “N” trace fluorescence under every non-“N” basecall (e.g., T fluorescence under A, C or G peaks) is compiled to generate a sample of the noise distribution. The sample of the noise distribution for each base is fitted to a zero-adjusted gamma distribution (zΓ, Supplementary Figure S2) using the package *gamlss* (23). We chose the zΓ distribution for three main reasons; 1) it has a domain from 0 to +∞, 2) it is a continuous distribution allowing for non-integer values, and 3) it allows for a high proportion of zeros in the data, which accounted for 10% of the values in our data (Supplementary Figure S2) (23). Filliben’s correlation coefficient (*R*_F_^2^) is calculated to assess the “goodness of fit” of the model given the data, where *R*_F_^2^ = 1 is a perfect fit. From this model we can assign critical values using a default level of significance (α = 0.01), which the user can manually change on EditR’s interface.

EditR was written in the R statistical programming environment (v. 3.4.0). EditR requires a sample AB1 Sanger sequencing file (i.e. cells treated with base editor and gRNA) and a 15-20 nt character string of the edited region of interest (i.e. gRNA protospacer). Initial parameters for the program have set defaults, or can be adjusted by the user under the advanced settings. The EditR web app was written with the R package *shiny* (v. 1.0.1) and helped by incorporating design from TIDE and Poly Peak Parser (16, 17). The former identifies simple indel mixtures from Sanger sequencing data, while the latter calculates the frequency and composition of complex indel mixtures.

The sample file is uploaded and read into EditR. The fluorescence area of all four bases at each base call is assigned, as measured by the software provided by the capillary electrophoretic instrument manufacturer and determined by the *makeBaseCalls* function of *sangerseqR.* The percent area of each base is calculated by dividing the total area of the focal base by the area of all the bases summed together. The guide sequence is then aligned to the primary sequence generated from the base calls using the ends-free overlap alignment algorithm in *pairwiseAlignment()* with *type = “overlap”* argument from the *Biostrings* package (24). Ends-free alignment was chosen as it aligned to a local match, while also being robust to changes in the first base of the guide, as well multiple base changes in the middle of the guide.

## RESULTS

### EditR Workflow

To analyze the mutation frequency, spectrum and significance of BE3 treated cells a 400 bp – 800 bp region encompassing the edited site is PCR amplified and sequenced by standard dideoxynucleotide chain-termination based capillary electrophoresis (Sanger method). DNA isolated from BE3 and gRNA treated cells with significant editing should demonstrate polymorphisms under C bases (antisense G) within the base editing window (~5 bp of the protospacer with BE3 for example, Figure 1a, 3). Generally, these base edits are C → T (antisense G → A), however there are several documented instances of non-canonical base editing (i.e. C → G or A) including our work here (2–4, 11).

**Figure 1.**
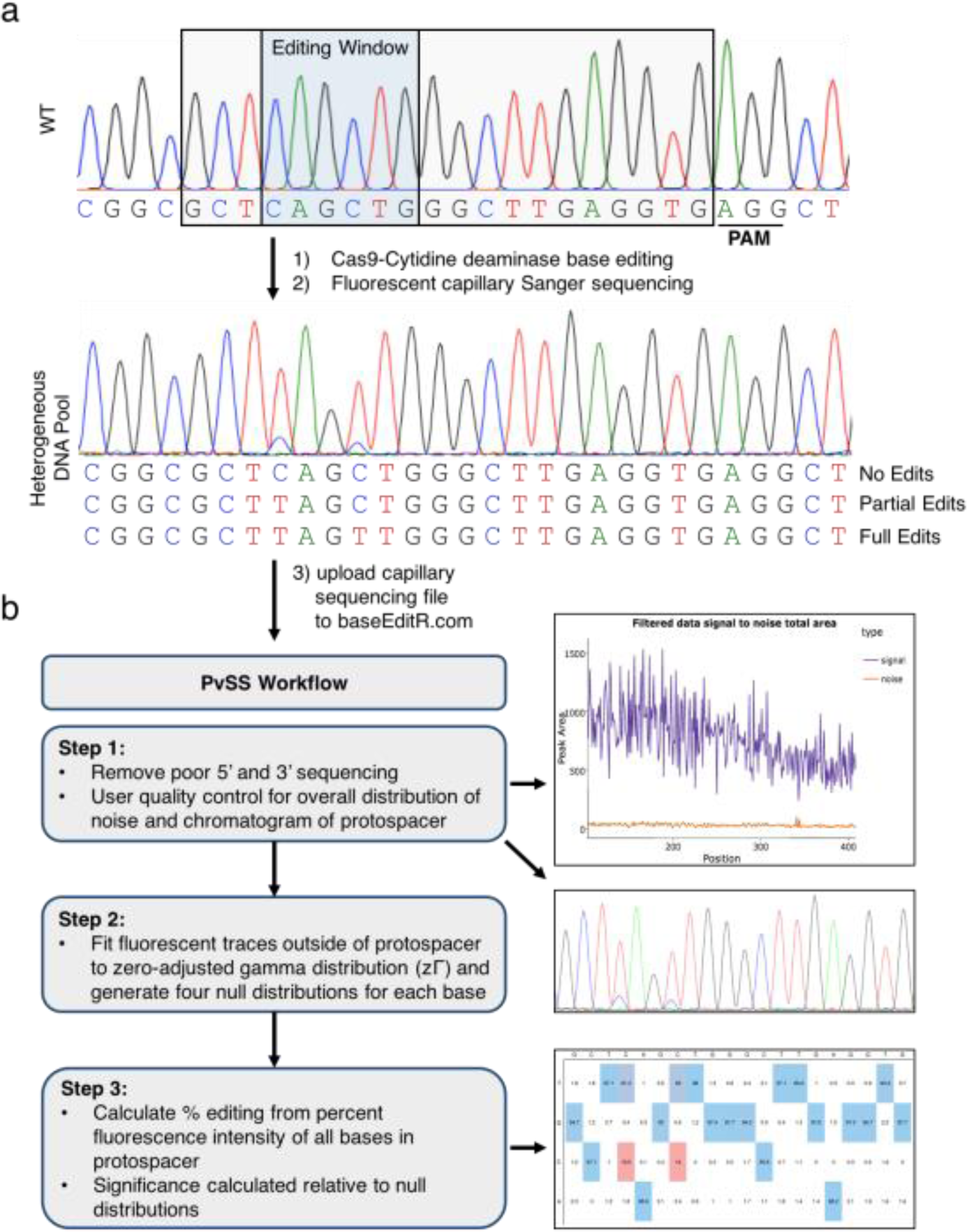
Analysis of base editing by capillary Sanger sequencing trace quantification. (**a**) Following treatment with base editor and gRNA, Cs (antisense Gs) within the editing window are converted to Ts (antisense As) producing a heterogeneous population of edited cells. (**b**) Workflow of EditR steps with summary plots: 1) The first and last portions of the file are removed due to poor quality. The signal-noise plot allows users to visualize the amount of fluorescence at each basecall that is deemed signal (purple) vs. noise (orange). A chromatogram of the protospacer is also produced for users to qualitatively validate their results. 2) A zero-adjusted gamma distribution is fit to the percent area noise of each trace (A,C,G and T) to generate four null distribution to which the traces in the hypothesized sites of editing (i.e. the protospacer) are compared. 3) Percent composition of traces measured to be significantly different from noise are plotted in a colored heat map proportional to magnitude (red is low, blue is high).

EditR generates a graphic of the percent noise across the sequencing file, allowing the user to assess the sequencing quality (Figure 1b, Step 1). If low quality regions are not filtered out by default settings, users can modify the region used to generate the null distribution. A chromatogram of the protospacer is generated to determine if the gRNA is properly aligned to the sequencing file and to visualize if the predicted editing matches qualitative expectations (Figure 1b, Step 1). The sequence traces within this region are compared to the traces in the rest of the sequencing file to quantify and determine the significance of base editing. EditR decomposes the trace at each basecall position into the percent fluorescence contribution of each of the four bases; A,C, G and T. The value of each percent “N” fluorescence at every “non-N” basecall is used to model a zΓ distribution, resulting in one zΓ distribution for each nucleotide. From these zΓ distributions a critical value is calculated as determined by the level of significance, which serves as the threshold for calling an edit within the protospacer as significant (Figure 1, Step 2, Supplementary Figure S2b). Percent editing is then calculated for traces within the protospacer that are above this threshold, the output of which is a heat-mapped table to visualize percent editing across the protospacer (Figure 1b, Step 3). The P_zΓ_-value in this context is the probability of calling a fluorescent peak a significant edit, when in fact that peak was merely noise rather than a base edit. On the EditR web app users can download a report of the results and a summary of the operations performed on their data.

### *In vitro* validation of EditR

To determine if quantitative sanger sequencing can accurately measure base editing under simulated conditions, we mixed together two PCR products that differed only in the presence of a single C vs. T. In two separate trials of different guide sites, samples were mixed in titrated amounts from 0 to 100% and subjected to capillary Sanger sequencing (Figure 2a, second trial Supplementary Figure S4). The calculated percent C→T agreed well with the actual concentration of PCR products by measuring either C or T in the two trials (*R*^2^ = 0.984). As a comparison to an alternative method of measuring base editing, titrations were also subjected to the Surveyor nuclease assay and quantified with fluorescence gel densitometry (Figure 2a, Supplementary Figure S3). The calculated percent editing as calculated by the surveyor assay agreed well with the actual concentration of the PCR products (*R*^2^ = 0.981), and with calculations by EditR showing that EditR is slightly more accurate than the surveyor assay in measuring base editing efficiency (Figure 2a).

**Figure 2.**
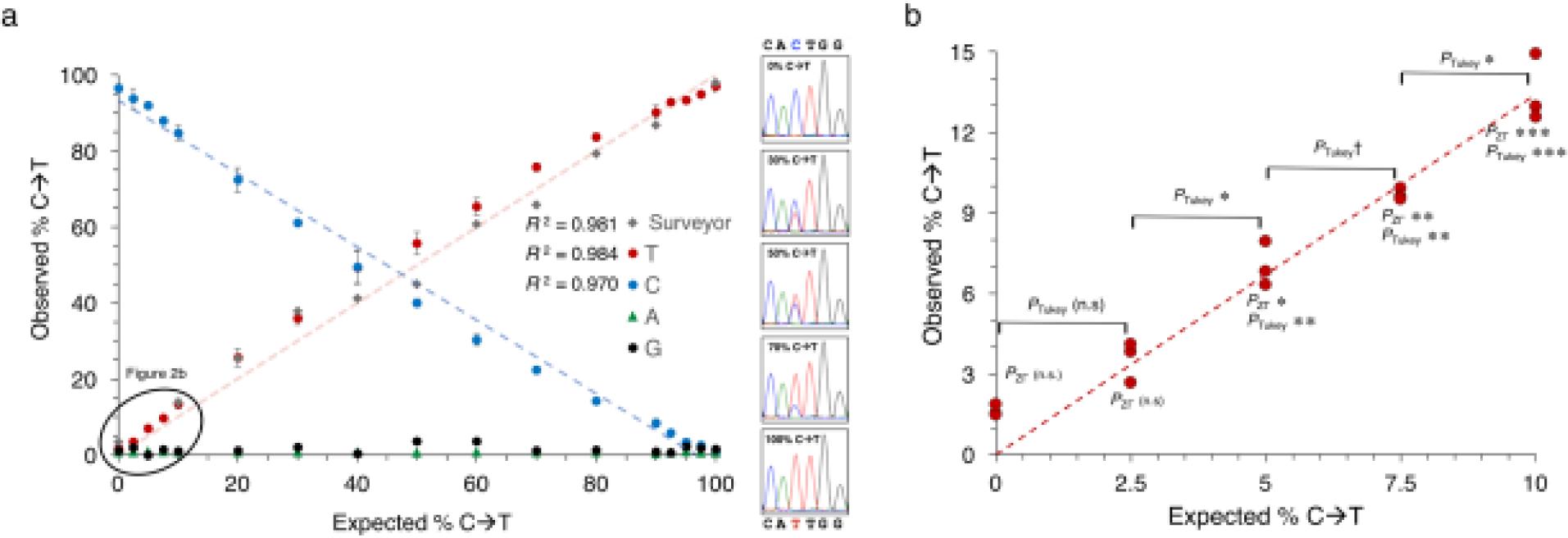
*In vitro* validation of EditR as an accurate and sensitive method. (**a**) EditR data presented as the mean of independent triplicates ± 1 standard deviation. See Supplementary Figure S4 for an additional titration series. Surveyor assay data presented as the mean of triplicate measurement ± 1 standard error of gel fluorescence densitometry as previously described (15). When the expected editing rate is above 50% the observed editing is calculated as 1 - (*calculated editing*) as the T containing product shifted from the minor to major PCR product. See Supplementary Figure S3 for gel image. (**b**) EditR can detect 2.5% differences in C→T down to 2.5% C→T. †*P*<0.05, \**P*<0.01, \*\**P*<0.001, \*\*\**P*<0.0001, n.s is non-significant. See Supplementary Figure S5 for data at 100% C→T end of titration for C traces.

To determine the precision and sensitivity of EditR we performed statistical tests between differing titrations. ANOVA with post hoc Tukey’s HSD test of each titration compared to the WT titration (0% C→T) showed that titrations could be measured as significantly different from WT as low as 2.5% C→T (*P* < 0.01, Figure 2b). One-way ANOVA, followed by Tukey’s HSD *post hoc* test demonstrated that triplicate samples could resolve incremental differences as small as 2.5% increments down to, but not past 2.5% C → T (*P <* 0.05, Figure 2b). By comparison, the EditR zΓ significance testing was able to resolve C → T editing from background noise down to 5.0% (*P* < 0.01, Figure 2b). These results were mirrored with the percent C area at the 95% C→T end of the spectrum (Supplementary Figure S3). This data demonstrates EditR is a sensitive and precise method for discerning and measuring low-level mutations likely to be observed in base edited cells.

### Application of EditR to Base Edited Cells with Canonical Mutations (C→T)

To assess the functionality of EditR in base edited cells, we treated HEK 293T cells with pCMV-BE3 and pENTR221-U6-gRNA. As expected, PCR amplification and capillary Sanger sequencing of the target site demonstrated noisy initial sequencing followed by a several hundred basepair span with a high percent signal (S/(S+N) ≥ 0.9) (Figure 3a). Informatively, the quality control plot generated by EditR showed two “noise” peaks within the highlighted protospacer region, which the editing quadplot confirmed to be from base editing of C→T (Figure 3b). Percent editing as calculated by EditR was consistent with percent editing as measured by the surveyor assay across three different targets, while in contrast to the surveyor assay, EditR was also able to distinguish the position and type of mutation (Figure 3c-h). Importantly, EditR was also able to determine the discrete editing efficiency in a multiply base edited sample (Figure 3c) and measure editing as low as 7.3% (*P*_zΓ_ < 0.01, Figure 3c,e). This data shows that EditR is an effective and reliable tool for measuring canonical C→T and G→A mutations in base edited cells.

**Figure 3.**
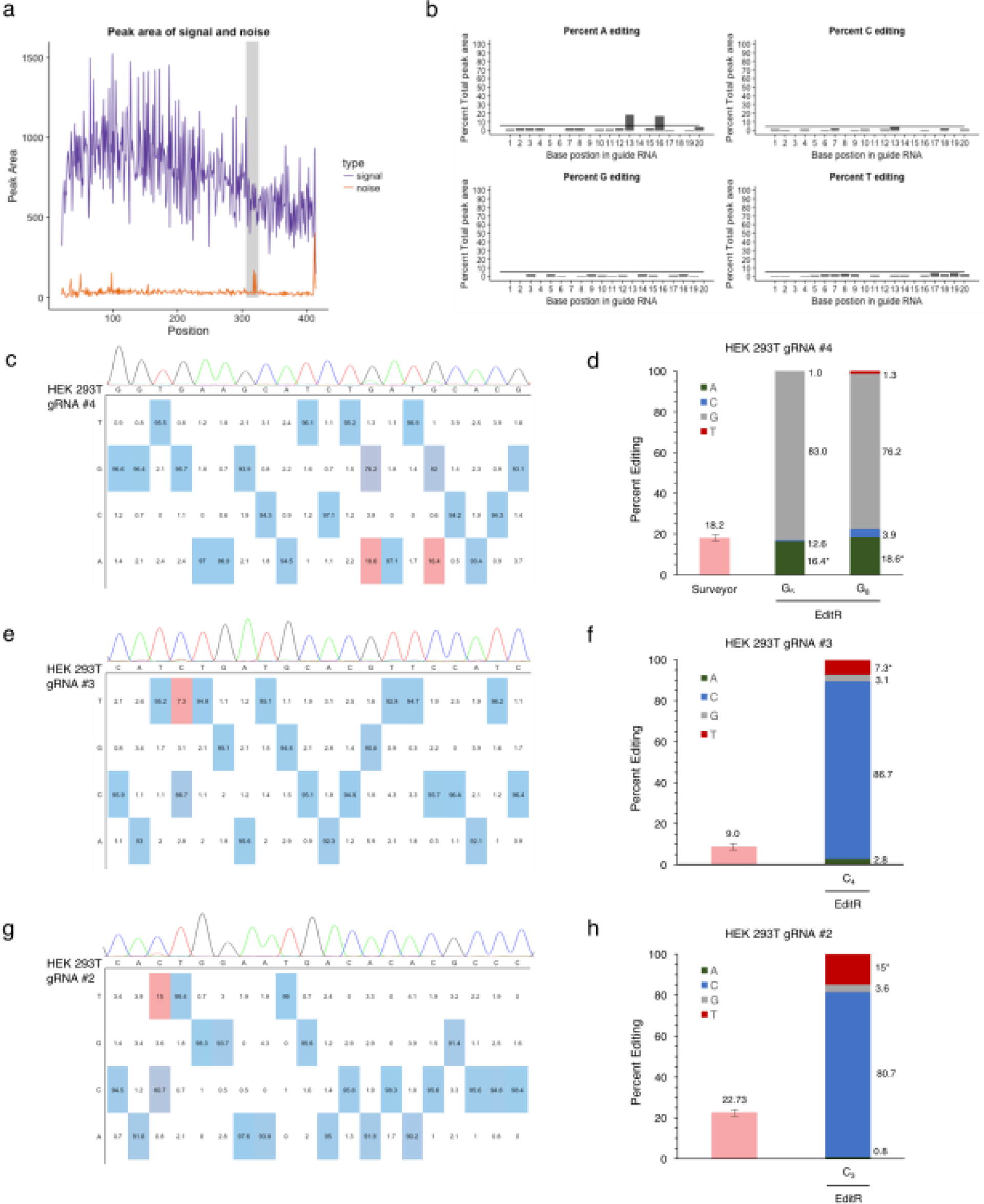
Validation of EditR in base edited cells with canonical mutations. (**a**) Output graphic showing the distribution of signal and noise in the sequencing file with peaks in the gRNA region. (**b**) Output plots of traces by base identity and significance. (**c**) Output editing table color-coded by proportionality showing editing of G→A at two bases. (**d**) Comparison of EditR to Surveyor assay, height of bars is the mean of triplicate measurement ± 1 standard error of gel fluorescence densitometry as previously described (15). WT negative control not shown (0% editing). See Supplementary Figure S3 for gel image. (**e-h**) Additional EditR generated table plots of base edited samples with canonical mutations.

### Application of EditR to Base Edited Cells with Non-Canonical Mutations (C→G or A)

To assess the functionality of EditR in measuring the frequency of non-canonical mutations which are regularly seen with base editors (1, 2, 6, 13, 14), we treated HOS and HCT116 cell lines with BE3 and gRNA using our previously published enrichment method that selects for highly edited cells (15). Sanger sequencing of cells treated with gRNA #1 was confirmed to be of high quality (S/(S+N) ≥ 97.5%, Figure 4a) and demonstrated ~40% base editing of Cs at positions 4 and 7, with C_4_ exhibiting a non-canonical C→G mutation and C_7_ exhibiting a canonical C→G mutation (Figure 4b,c). The surveyor assay yielded an editing efficiency of 39.1% which was similar to the 39.4% of C_4_ and 38.8% of C_7_. Because the percent G at C_4_ was nearly identical to the percent C at C_7_, it is suggestive that C_4_-T_7_ are linked together and account for 40% of the allelic pool, while G_4_-C_7_ are linked accounting for the 60% remainder. Further use of EditR shows its ability to resolve complex mixtures of non-canonical mutations in base edited cells across multiple cell lines and target sites. (Figures 4e-h). This demonstrates EditR can measure the editing efficiency of non-canonical mutations, while having the advantage over the surveyor assay in elucidating the discrete composition of non-canonical mutations.

**Figure 4.**
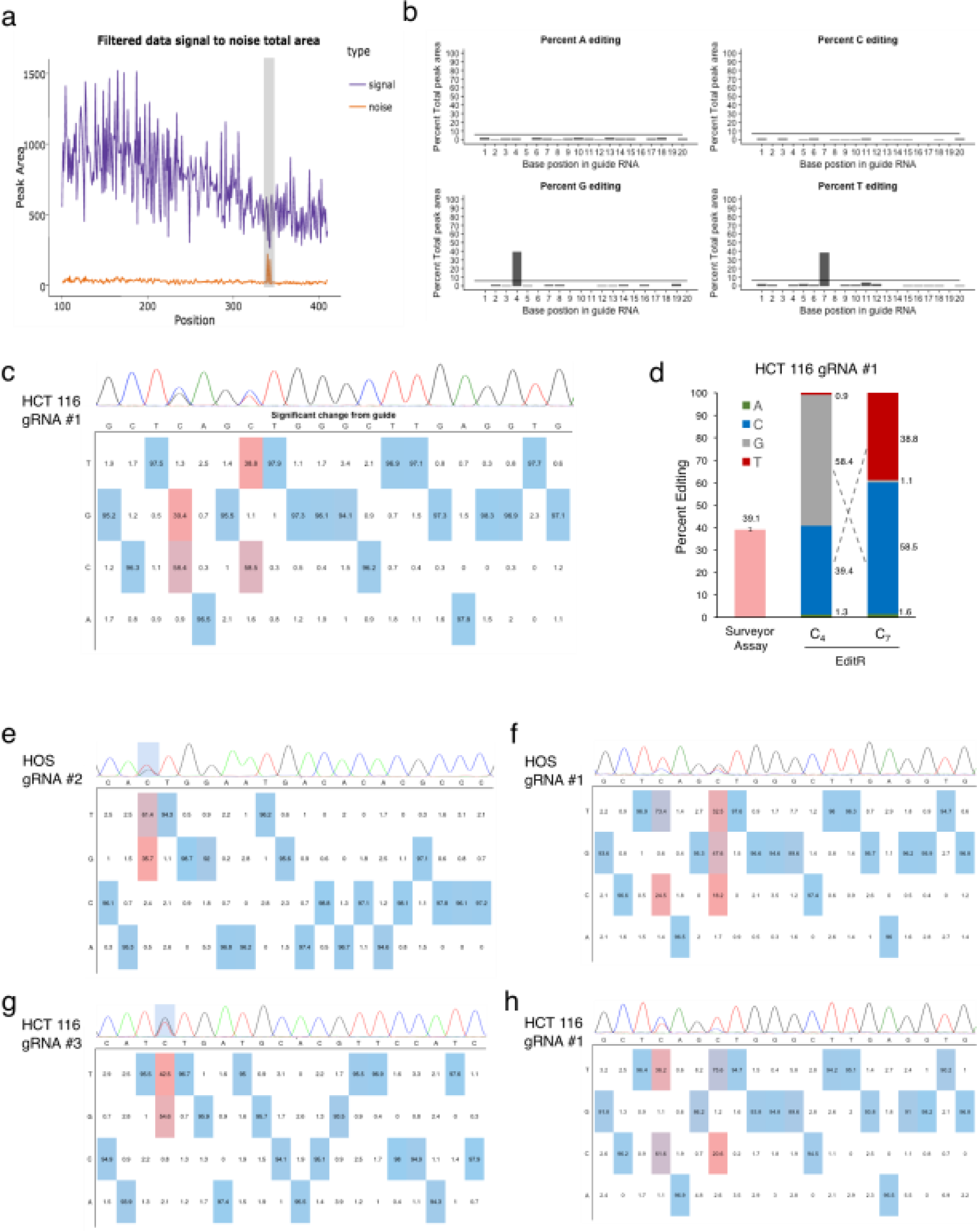
Validation of EditR in base edited cells with non-canonical mutations. (**a**) Output graphic showing the distribution of signal and noise in the sequencing file with peaks in the gRNA region (**b**) Output plots of traces by base identity and significance. (**c**) Output editing table color-coded by proportionality showing significant C→T and C→G mutations. (**d**) Comparison of EditR to Surveyor assay, height of bars is the mean ± 1 standard deviation of fluorescence densitometry from independent surveyor assays. Similarity in the percent editing suggests linked mutations. See Supplementary Figure S3 for gel image. (**e-h**) Additional EditR generated table plots of base edited samples with canonical mutations.

### Comparison of EditR to Next-Generation Deep Sequencing

To assess potential tradeoffs of the ease of using EditR against the accuracy of its measurements, and to assess the accuracy of EditR in multiple sequence contexts, we compared EditR to next-generation deep sequencing (NGS) which is the gold-standard for measuring base editing (3, 4, 10, 11). HEK 293T cells were treated with BE3 and one of nine gRNAs that targeted one of five genomic sites (Figure 5a, Supplementary Figure 1). Genomic DNA was harvested from treated cells, PCR amplified for the edited region of interest, and amplicons were concurrently Sanger sequenced and deep-sequenced to directly compare EditR to NGS. ΕditR yielded measurements of base editing that were not significantly different from NGS by paired t-test (*P = 0.27, df = 27,* Figure 5b,d), with an average difference of 0.74% (99% CI −0.62% to 2.11%, Figure 5c,d) and standard deviation of 3.45%. Furthermore, samples were confirmed by NGS to possess non-canonical mutations spanning the spectrum of C→T, A or G (Supplementary Figure S6). The lack of a significant difference between measuring base editing by EditR vs. NGS shows that EditR is a highly robust method of measuring canonical and non-canonical base editing outcomes in multiple sequence contexts.

**Figure 5.**
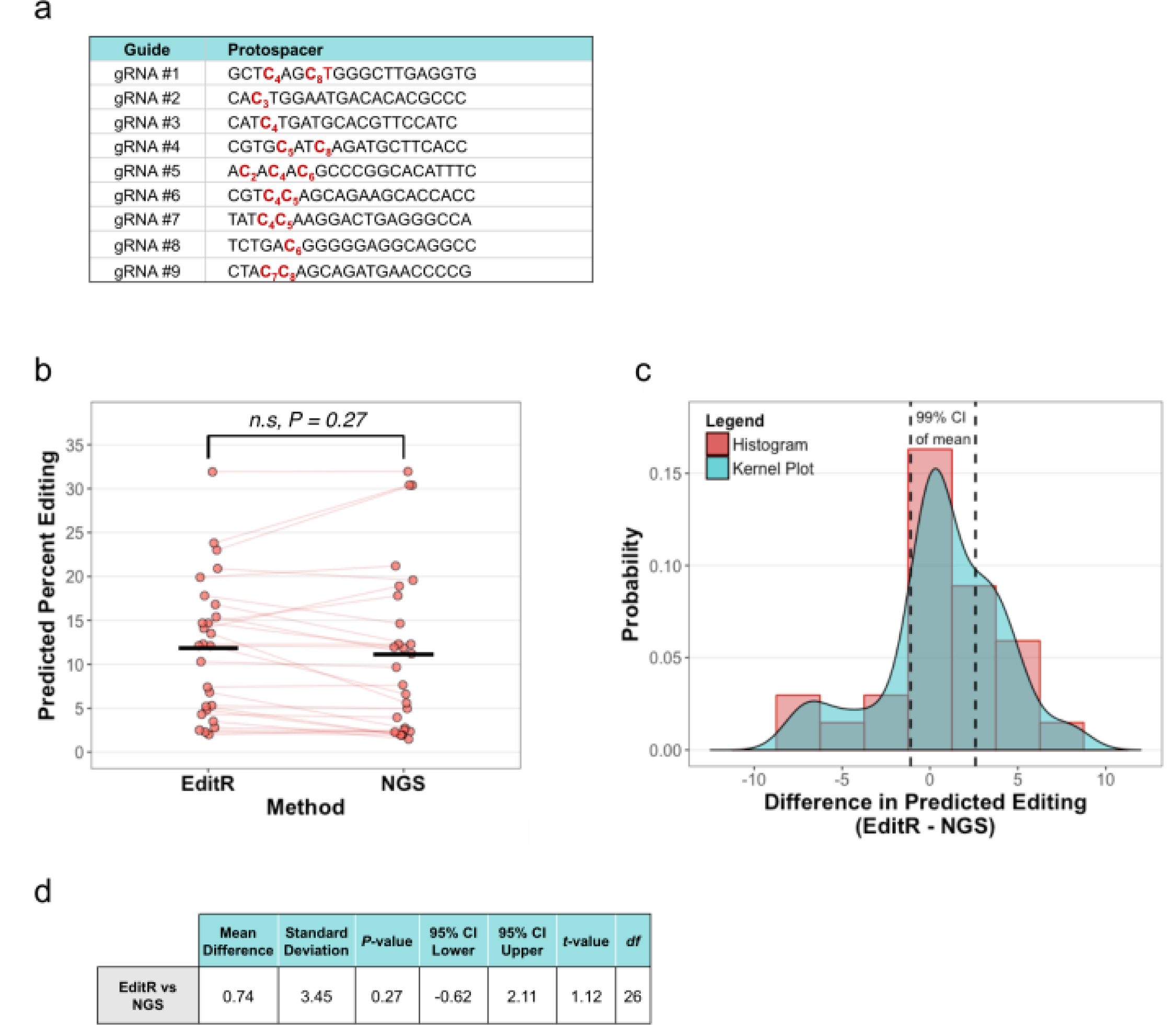
Validation of EditR compared to next-generation deep sequencing (NGS) across multiple guide sites. (**a**) Base editing guides used to compare EditR to NGS. (**b**) Comparison of measured editing by EditR and NGS, solid black bars denote the mean of each group, red lines drawn between points indicate which samples are paired. (**c**) Distribution of differences between EditR and NGS, positive values indicate editing measured by EditR was larger. (**d**) Table summary of paired t-test performed on data. N = 27, comprised of 5 amplicons, 9 unique guides, 17 unique bases, and 21 unique edits with 1 or 2 independent replicates per guide. A single unique base may produce multiple unique edits via non-canonical editing, i.e., one unique C edited to T, G or A.

## DISCUSSION

### Advantages of EditR

Cas9-Cytidine deaminase base editors are a new but rapidly expanding technology, with potential applications spanning the biomedical sciences. The versatility of base editing is astronomical (1, 10), requiring an equally adaptable method to analyze base editing outcomes. Here we show that the Surveyor nuclease assay can accurately measure base editing mutations, however it is unable to resolve the composition and position of base editing. While there are several other methods available to measure base editing efficiency, all suffer from poor accuracy or high costs hindering access to base editing research. Given the high requirement of resources needed to accurately measure base editing this creates an accessibility barrier in base editing research which disfavors genuine competition and collaboration. As an alternative, we developed EditR as a rapid, accurate and inexpensive approach to measuring base editing efficiency. EditR takes advantage of the proportional change in the percent area of trace fluorescence as bases are edited. This percent area is compared to the background distribution of percent fluorescence noise to determine if significant editing is occurring. EditR enables researchers to both quantify base editing by position and to assess the composition of mutations at a particular base. Comparatively, EditR is a fraction of the cost of next generation deep sequencing with returned results possible within a day. This makes EditR a useful alternative to, or as a pre-sequencing validator of, NGS.

### Comparison to other programs for base editing research

The resolution and accuracy of EditR is equal to that of other programs that quantify nucleotide polymorphisms from Sanger sequencing such as QSVanalyser (http://dna.leeds.ac.uk/qsv/) and Mutation Surveyor^®^ (http://www.softgenetics.com/mutationSurveyor.php) (>5% resolution) (25, 26). While these programs are highly useful for analyzing discrete single-nucleotide polymorphisms (SNP) or copy number variants they are less suitable for base editing research. The algorithms of Mutation Surveyor and QSVanalyzer both rely on adjacent peaks as a reference to the base of interest when measuring editing efficiency (25, 26). For example, QSVanalyzer compares the intensity of the base of interest to the heights of the peaks between 5 and 10 bases upstream of the base of interest to measure the percentage of the minor SNP (25). This referencing method is powerful when looking at discrete single point mutations, but is less amenable to base editing, as base editors are processive enzyme that will edit adjacent cytidines within the editing window. This issue is especially relevant when considering new generations of base editors, some of which have editing windows as large as 14 nucleotides (10). EditR overcomes these issues by comparing the trace within the protospacer against the background distribution of noise outside of the protospacer instead of adjacent peaks. Furthermore, EditR is accessible and intuitive as a free web application, or as open source code that can be ran locally as an R-shiny app on any major operating system.

### Limitations of EditR

EditR is largely limited by the quality of the Sanger sequencing results, because EditR measures base editing by determining if trace fluorescence is due to editing or noise. To account for this, we recommend gel extracting or purifying PCR products with a proprietary kit prior to sequencing. We advise using traces that have an average percent noise of ≤ 7.25% and modelled parameter μ of ≤ 2.5 as that is strongly correlated with EditR calling significance at *P < 0.01* (Supplementary Figure S7a-b). Furthermore, it is important that the zΓ models are properly fit in order to have sensitive detection of base editing, thus we recommend only using sequencing files with an R_F_^2^ ≥ 0.95 as we found the vast majority of our chromatograms fall in this range (Supplementary Figure S7c). As a note, even in files with a large proportion of noise, R_F_^2^ was still above 0.9 showing that even in noisy samples the zΓ distribution effectively models the noise distribution (Supplementary Figure S7d).

In considering the precision of EditR, our results indicate that ~95% of samples analyzed using EditR will deviate by no more than −6.1% to +7.5% (mean ± 2SD; ± ~6.8%) from the percent editing as measured by NGS (source). The precision of EditR is similar to that of TIDE (mean ± 2D = −5.4% to +4.2%, Figure S8), which further supports a quantitative analysis of sanger sequencing to analyze gene editing efficiency. To assess what may cause EditR to deviate from NGS values, future work needs to address how local sequence contexts alter percent fluorescence area. In fluorescent Sanger sequencing the identity of the preceding base affects the intensity of the subsequent base, but it is not clear how certain sequence contexts my affect calculations of editing efficiency (25). For example, EditR may be unable to accurately measure some types of base editing in certain sequence contexts such as repetitive G-rich reads (25, 27). As such, future work will develop algorithms that incorporate the local sequence context to adjust the predicted editing efficiency to more accurately quantity base editing.

### Future Applications of EditR

Here, we used the base editor BE3 as the basis of our work, however EditR could be used with other base editors that target outside of the protospacer (3, 4) by having the user adjust the edited region of interest. EditR could also be used to measure rates of Cas9 mediated homology directed repair, where a small to moderate region of a gene is replaced which could be set as the edited region of interest. In general, any research concerned with measuring rates of localized nucleotide variation could use the EditR method. As such, EditR’s algorithm could be adapted for quantitative bisulfite sequencing by excluding CpG dinucleotides from the null sample and then measuring the traces at CpG islands to see if they are significant relative to the null distribution. Ultimately, the EditR approach has many potential applications within and outside of genome engineering, and in tandem with TIDE (16), these approaches represent the ability to affordably and accurately measure nearly any outcome of contemporary Cas9 mediated genome editing.

Within the past year of base editing research, there has been extensive modification to the core enzymatic machinery, including interchange of the Cas9 and APOBEC domains for improved versatility or precision (10) and the additional fusion of the ssDNA binding protein Mu GAM to reduce non-canonical mutations (11). With innovation in mind, it is likely that other mutagenic enzymes will also be constructed in a modular fashion to expand the directionality of base editing beyond C→T mutations. A likely candidate for the next generation of base editors is an Adenosine Deaminase (ADAR) fusion enzyme. ADAR deaminates adenosine to inosine (A→I) in RNA *in vivo* with preliminary evidence of dsDNA A→I deamination *in vivo* (28). In dsDNA inosine behaves predominantly like guanine forming I:C base pairs, resulting in an A→G mutation (29). Recently, and RNA guided ADAR system was leveraged to edit the ssDNA M13 bacteriophage genome *in vitro* (30) suggesting ADAR mediated base editing of genomic dsDNA is feasible. With the programmable specificity of Cas9 fused to the directional mutagenicity of ADAR, it is highly probably an A→G (antisense T→C) base editor is on the horizon. No matter the directionality of the next generation of base editors, EditR is equipped to analyze the future of this burgeoning field.

## AVAILABILITY

baseEditR.com/

https://github.com/MoriarityLab/EditR

## SUPPLEMENTARY DATA

Supplementary Data are available on NAR Online.

## ACKNOWLEDGEMENT

M.G.K and B.S.M designed the research. M.G.K and W.S.L constructed all vectors. M.G.K, W.S.L and B.R.W performed all transfections and generated all clones. M.G.K performed all Surveyor nuclease assays. M.G.K and D.A.N wrote the program. D.A.N. designed the online application. M.G.K, D.A.N. and B.S.M wrote the article. J.R.G and J.E.A performed bioinformatics analyses. Thank you to Arianna Wegley for the helpful suggestions on the manuscript, Leah Hogdal for beta testing the web application, and Yaniv Brandvain for conversations around developing the statistical testing approach.

